# A high-throughput zebrafish screen identifies novel candidate treatments for Kaposiform Lymphangiomatosis (KLA)

**DOI:** 10.1101/2024.03.21.586124

**Authors:** Ivan Bassi, Amani Jabali, Naama Farag, Shany Egozi, Noga Moshe, Gil S. Leichner, Polina Geva, Lotan Levin, Aviv Barzilai, Camila Avivi, Jonathan Long, Jason J. Otterstrom, Yael Paran, Haim Barr, Karina Yaniv, Shoshana Greenberger

## Abstract

Kaposiform Lymphangiomatosis (KLA) is a rare, aggressive, and incurable disease caused by a somatic activating NRAS mutation (p.Q61R) in lymphatic endothelial cells (LECs). The development of new therapeutic avenues is hampered by the lack of animal models faithfully replicating the clinical manifestations of KLA. Here, we established a novel zebrafish model of KLA by driving conditional expression of the human NRAS mutation in venous and lymphatic ECs. We find that mutant embryos recapitulated clinical features of KLA, including pericardial edema and a dilated thoracic duct, and that the phenotypes were reverted by Trametinib, a MEK inhibitor used for KLA treatment. We further leverage this model in combination with an AI-based high-throughput drug screening platform to search for small compounds selectively reverting the mutant phenotypes and identify Cabozantinib, an FDA-approved tyrosine kinase inhibitor, and GSK690693, a competitive pan-Akt kinase inhibitor, as leading hits. Finally, we test these drugs in cultured cells derived from KLA patient and demonstrate their ability to normalize LEC sprouting and block NRAS downstream pathways, underscoring the potential of GSK690693 and Cabozantinib as potential KLA treatments. Overall, our novel zebrafish model provides a valuable tool for research into the etiology of KLA and for identifying new therapeutic avenues.

## Introduction

Kaposiform lymphangiomatosis (KLA) is a rare, infiltrative, multifocal, or diffused lymphatic malformation classified as a new subtype of generalized lymphatic anomaly (GLA)^1^. The vast majority of diagnoses are made in children, with a mean age of 8.2 ^2^, but rare reports of KLA have also been described in adults, with a similar presentation ^3^. Clinical manifestations of KLA include subcutaneous and mediastinal masses, bleeding, and pleural effusions. The mortality rate from KLA is high, with cardiorespiratory failure as the leading cause of death among patients ^4^. Current pharmaceutical treatments for KLA, such as rapamycin (sirolimus)^5^ and MEK inhibitors ^6^ have led to clinical improvement and potentially increasing survival rates ^7^. Nevertheless, the response to these treatments varies among patients, and the overall survival rate for KLA remains disappointingly low. Moreover, there is still no standard therapeutic approach ^4^, underscoring the critical need for effective medical interventions to manage KLA.

Our previous work identified for the first time a somatic activating NRAS mutation (p.Q61R) in lymphatic endothelial cells (LECs) from involved tissues of a GLA/KLA patient ^8^. The Q61R mutation results in an amino acid substitution at position 61, from glutamine (Q) to arginine (R). This hotspot mutation lies within the GTP-binding region of the NRAS protein and results in increased GTP-bound NRAS ^9,10^. This subsequently leads to constitutive activation of the RAS-MAPK and PI3K-AKT-mTOR downstream pathways^11^ promoting cell growth, survival, and invasion. Our original results have been further replicated *in vitro*^12^ and genomic evaluation of KLA patients conducted in recent years further identified the Ras pathway, particularly NRAS p.Q61R, as the likely driver of the disease in most cases^13,14^.

The progress in understanding the molecular mechanisms underlying KLA pathogenesis and the subsequent development of new therapeutic avenues is hampered by the lack of reliable animal models capable of faithfully replicating the diverse clinical manifestations observed in KLA. Our previous work demonstrated the abnormal development of the zebrafish blood and lymphatic vasculature following transient pan-endothelial overexpression of the mutant form of human NRAS, highlighting the potential of zebrafish as an advantageous model to study KLA^8^. Owing to their rapid external development, optical transparency, high number of offspring, and straightforward strategies for genetic manipulations, zebrafish provide excellent means for modeling human diseases. In addition, by combining the scale and throughput of *in vitro* screens with the physiological complexity of animal studies, the zebrafish has already contributed to several successful phenotype-based drug discoveries^15^. Recently, for instance, a lifesaving treatment for a lymphatic anomaly was identified through such a high-throughput drug screen conducted in zebrafish embryos carrying a human mutation^16,17^, highlighting the potential of zebrafish-based studies for discerning disease molecular mechanisms and translating them to the clinics.

The zebrafish possesses a lymphatic system that shares many characteristics of lymphatic vessels found in other vertebrates^18^, including the requirement for the Prox1 transcription factor for proper lymphatic formation^19,20^ and the expression of *lyve1b* the zebrafish orthologue of *Lyve1*, the gold standard marker for human lymphatics^21–23^. Live imaging of zebrafish embryos established that the parachordal cells (PACs), which form along the embryo’s midline at 2 days post-fertilization (dpf), serve as building blocks for the lymphatic system^18,19^. Starting at 2.5 dpf, PACs migrate ventrally to generate the main lymphatic vessel, the thoracic duct (TD), which runs along the fish trunk between the Doral Aorta (DA) and the Cardinal Vein (CV), resembling its anatomical location in mammals.

Here, we describe the establishment of a novel zebrafish model of KLA by driving conditional expression of the mutant form (p.Q61R) of human NRAS (*hNRASmut*) in venous and lymphatic ECs. We show that induction of *hNRASmut* expression in zebrafish embryos at 2 dpf results in pericardial edema and a markedly dilated TD, recapitulating some of the clinical manifestations of human KLA. Moreover, we demonstrate that treatment with the MEK inhibitor Trametinib, currently used in the clinic, effectively rescues all the phenotypes associated with the mutation, thereby confirming the suitability of our model. We further leverage this system to screen for small compounds selectively reverting the mutant phenotypes and identify Cabozantinib, an FDA-approved tyrosine kinase inhibitor, and GSK690693, a competitive pan-Akt kinase inhibitor, as leading hits. Finally, we test these drugs in cultured cells derived from KLA patient and demonstrate their ability to normalize LEC sprouting and block NRAS downstream pathways, underscoring the potential of GSK690693 and Cabozantinib as novel therapeutic avenues for treating KLA.

## Results

To establish a conditional zebrafish model of KLA, we first expressed the wt and mutant forms of human NRAS under the UAS promoter (Fig. S1A). The newly generated Tg*(UAS:hNRASp.Q61R)* fish (hereafter *hNRASmut*) were initially mated with Tg*BAC(prox1a:KALTA4;4xUAS- E1B:TagRFP)*^24,25^, to drive constitutive expression of the *UAS:NRAS* wt and mutant constructs in LECs. At 5 dpf *prox1a:KALTA4;hNRASmut larvae* exhibited profound defects, such as swollen bodies and pronounced pericardial edema (Fig. S1B,C red arrow). Confocal images also revealed a severely malformed vasculature, with a markedly dilated posterior cardinal vein (PCV) as well as collapsed and missing intersegmental vessels (ISVs) (Fig S1D,E). Given the broad and early embryonic expression of the *prox1a* gene, which also includes neurons and somites, among other tissues^26^, we suspected that part of these phenotypes could result from overall toxicity due to ectopic NRAS expression. To overcome this, we generated a conditional and inducible model of KLA by crossing Tg*(UAS:hNRAS)* wt and mutant animals with fish expressing the Gal4 transcriptional activator under the regulation of the venous and LEC-specific promoter *lyve1b*^22–24^. To avoid early lethality and /or toxicity, we fused the *Gal4* construct to the estrogen receptor variant (ERT2) (hereafter *Tg(lyve1b:Gal4^ERT^*^2^). Upon the administration of 4-hydroxytamoxifen, ERT2 is rapidly activated, thus inducing expression of the UAS constructs^25,27^. As seen in Fig. 1A-B’ targeted expression of *hNRASwt* in venous and lymphatic ECs at 24 hours post fertilization (hpf) did not cause any noticeable defects at 5 dpf. In contrast, the presence of *hNRASmut* in the same EC population resulted in severely swollen embryos with pronounced pericardial edema (Fig. 1C,C’, red dashed line). Measurement of the pericardial area revealed a 4-time increase in *hNRASmut* embryos, as compared to their wt and *hNRASwt* expressing counterparts (Fig. 1D). Interestingly, we noticed that activation of *hNRASmut* at 48 hpf does not affect the normal development of the embryos or leads to changes in overall body morphology or pericardial edema, as observed following induction at 24 hpf (Fig. S1F).

**Figure 1.**
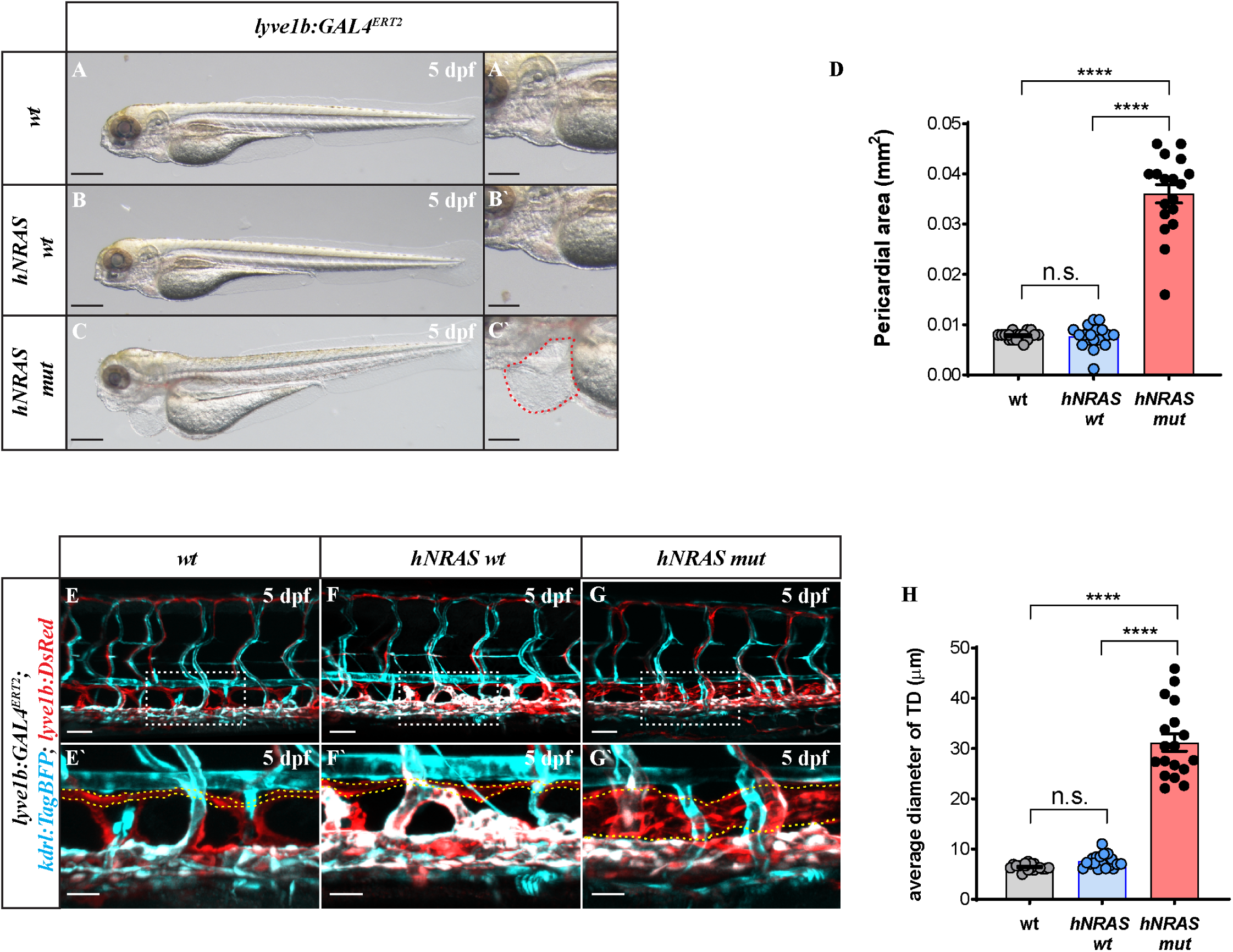
Establishment and characterization of a zebrafish KLA model. **A-C’.** Phase Contrast images of Tg(*lyve1b:Gal^ERT2^*) wt fish (A,A’), overexpressing h-NRAS wt (*hNRASwt*, B,B’) or hNRAS p.Q61R (*hNRASmut*) (C,C’) variants. Only mutant *larvae* depict swollen bodies (C) and pericardial edema (C’, red dashed line). **D.** Quantification of pericardial area (n=18; One-way ANOVA, multiple comparisons with Tukey post-hoc test). **E-G’.** Confocal images of trunk vasculature showing enlarged lymphatic thoracic duct (TD) after induction of *hNRASmut* (G, G’); yellow dashed lines in E’-G’ delineate TD, dashed squares in E-G mark the area enlarged in E**’-**G’). **H.** Quantification of TD diameter in wt, *hNRASwt* and *hNRASmut larvae* (n=18; One-way ANOVA, multiple comparisons with Tukey post-hoc test). Scale bars: A,B,C = 100 μm; A’,B’,C’ = 50 μm; E,F,G = 50 μm; E’,F’,G = 50 μm. Error bars are mean ± s.e.m.

We next investigated the effects of *hNRASmut* expression on the developing vasculature by mating *Tg(lyve1b:Gal4^ERT2^;UAS:hNRASwt)* and *Tg(lyve1b:Gal4^ERT2^;UAS:hNRASmut)* fish with a double fluorescent reporter highlighting lymphatic (Tg(*lyve1b:dsRed))* and blood (Tg*(kdrl:TagBFP*)) vessels. Confocal images revealed that expression of *hNRASmut* but not *hNRASwt* in venous and lymphatic ECs leads to an abnormal, dilated TD at 5 dpf (Fig. 1E-H, TD marked in yellow). In contrast, we did not notice any defects in the caliber and morphology of the DA and the PCV (Fig. S1G,H), confirming that the expression of mutant NRAS affects specifically the lymphatic endothelium. Notably, we observed a complete correlation between pericardial edema and dilated TD phenotypes, as all embryos expressing the mutation displayed both conditions.

In zebrafish, the TD forms following migration and coalescence of sprouts from the PACs)- lymphatic progenitors situated at the level of the horizontal myoseptum, which serve as the building blocks of the fish lymphatic system^19,28^. To get insight into the stages of TD formation affected by the mutation, we first assessed the number of PACs at 3.5 dpf. As seen in Fig. S1I-K, no significant differences were detected when comparing wt and *hNRASmut* larvae, suggesting that the mutation does not affect the ability of LEC progenitors to sprout from the CV and reach the HM. Next, we imaged *Tg(lyve1b:DsRed);hNRASmut/wt* in time-lapse between 60-120 hpf, following induction at 24 hpf (Video S1). Starting at ∼60 hpf, PAC sprouts begin migrating dorsally and ventrally from their location at the HM, following the ISVs to establish the dorsal longitudinal lymphatic vessel (DLLV)^29^ and the TD just ventral to the DA^19^. As seen in snapshots extracted from the time-lapse series, we could detect the forming TD in wt larvae at ∼75 hpf (Fig. S1L’). At this stage, however, the TD appears already slightly enlarged in *hNRASmut larvae* (Fig. S1M’), and so do the emerging PAC sprouts (Fig. S1M’’, blue arrowhead). This phenotype becomes more pronounced as development proceeds (Fig. S1L’’, M’’). Interestingly, at ∼90hpf, we also noticed that *hNRASmut larvae* fail to establish a complete DLLV (Fig S1M’’, white asterisks), which appears fully established at this stage in wt siblings (Fig S1L’’, white arrowheads). Taken together, these results suggest that induction of *hNRASp.Q61R* mutation in the CV-ECs at 24 hpf does not affect the specification or differentiation of LEC progenitors but rather their subsequent proliferation, migration, and/or cellular morphology.

Recent studies have reported the positive effect of MEK inhibitors on complex lymphatic anomalies (CLA) in humans and zebrafish^17,30^. Therefore, we asked whether Trametinib treatment effectively reverted the lymphatic phenotypes in *hNRASmut* mutant *larvae*. Following induction of *hNRASmut/wt* expression at 24 hpf, we added Trametinib to the embryo water 24 hours later and carried out morphological analyses at 5 dpf (Fig. 2A). As seen in Fig. 2B-F, Trametinib treatment led to a complete recovery of the swollen body and pericardial edema phenotypes in *hNRASmut larvae* without affecting the normal development of wt embryos (Fig. 2C,C’,F). In addition, confocal images of Tg*(lyve1b:dsRed;kdrl:TagBFP) larvae* showed a significant rescue of the TD dilation upon Trametinib treatment (Fig. 2G-J’, TD marked in yellow, quantified in K).

**Figure 2.**
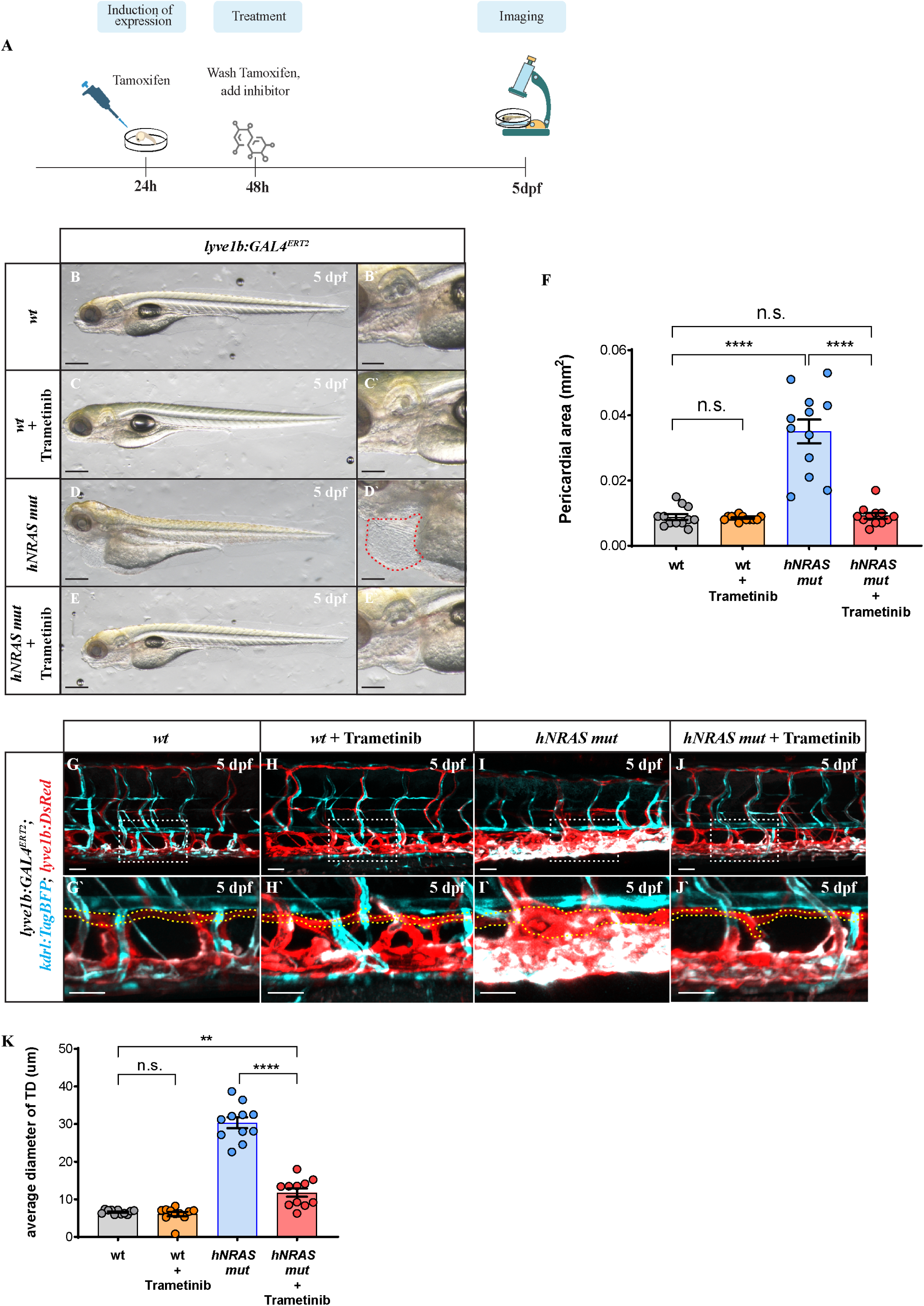
The MEK inhibitor Trametinib rescues the *hNRASmut*-induced phenotypes. **A.** Diagram illustrating the experimental design for drug testing. **B-E’.** Phase Contrast images of wt (B-C’) and *hNRASmut larvae* (D-E’), untreated (C,C’,E,E’) or following addition of Trametinib to the water (C,C’,D, D’, red dashed line in D’ depicts enlarged pericardial area). **F.** Pericardial area measurement indicates a complete recovery in *hNRASmut* embryos after Trametinib treatment (n=11; One-way ANOVA, multiple comparisons with Tukey post-hoc test). **G-K.** Confocal images (G-J’) of trunk vasculature showing enlarged TD after induction of *hNRASmut* (I,I’) that is reverted following Trametinib treatment (J,J’). Trametinib addition does not impact the development of wt embryos (C,C’). Yellow lines in G’-J’ delineate TD, dashed squares in G-J mark the area enlarged in G’-J’). Quantification of TD average diameter in wt, wt + Trametinib, *hNRASmut,* and *hNRASmut +* Trametinib *larvae* (K). Scale bars: B,C,D,E = 100 μm; B’,C’,D’,E’ = 50 μm; G,H,I,J = 50 μm; G’,H’,I’,J’ = 100 μm; Error bars are mean ± s.e.m.

Overall, the phenotypic traits observed in zebrafish *hNRASmut larvae*, such as dilated lymphatic vessels and pericardial edema, alongside the positive response to Trametinib treatment, mirror characteristics of KLA found in human patients, strongly supporting the validity of our model for the study of KLA.

An important factor underlying the lack of specific treatments for CLAs in general, and KLA in particular, is the lack of suitable animal models enabling efficient testing of potential therapeutic agents. We thus sought to take advantage of the easily distinguishable phenotypes of our newly generated *hNRASmut* zebrafish model to identify safe and effective treatments for KLA. To this end, we devised a high-throughput drug screening platform that relies on high-content imaging, coupled with a deep learning-based AI algorithm designed to analyze brightfield images of whole zebrafish *larvae* (WiSoft^®^ Athena software, see Materials and Methods and^31^ for details). We initially utilized 5 dpf *hNRASwt* and *hNRASmut* larvae, plus or minus Trametinib, to set up the system and adjust the previously trained AI algorithm^31^. Trametinib-treated *larvae* were regarded as a positive benchmark for "rescue," whereas untreated or DMSO-treated *larvae* served as negative controls (Fig. 3A-D). Only side-oriented fish, defined as those with an eye count and a tail count equal to one, were used for automated image analysis. Images were initially analyzed using AI-based segmentation to define estimated masks for the larva’s outer contour and three body compartments (head, trunk, and tail). Later, the masks were edited manually by adjusting their anchor points or using drawing tools to extend them. The manually annotated masks demarking the fish outline (Fig. 3A-D, red contour) were used to re-train the AI algorithm and create an improved detection model to quantify phenotypes. Data from our pilot experiments highlighted the significance of the "total larva area" (Fig. 3E) as the most effective measure for distinguishing between phenotypically wt and mutant animals. We thus applied this parameter to analyze all subsequent experiments.

**Figure 3.**
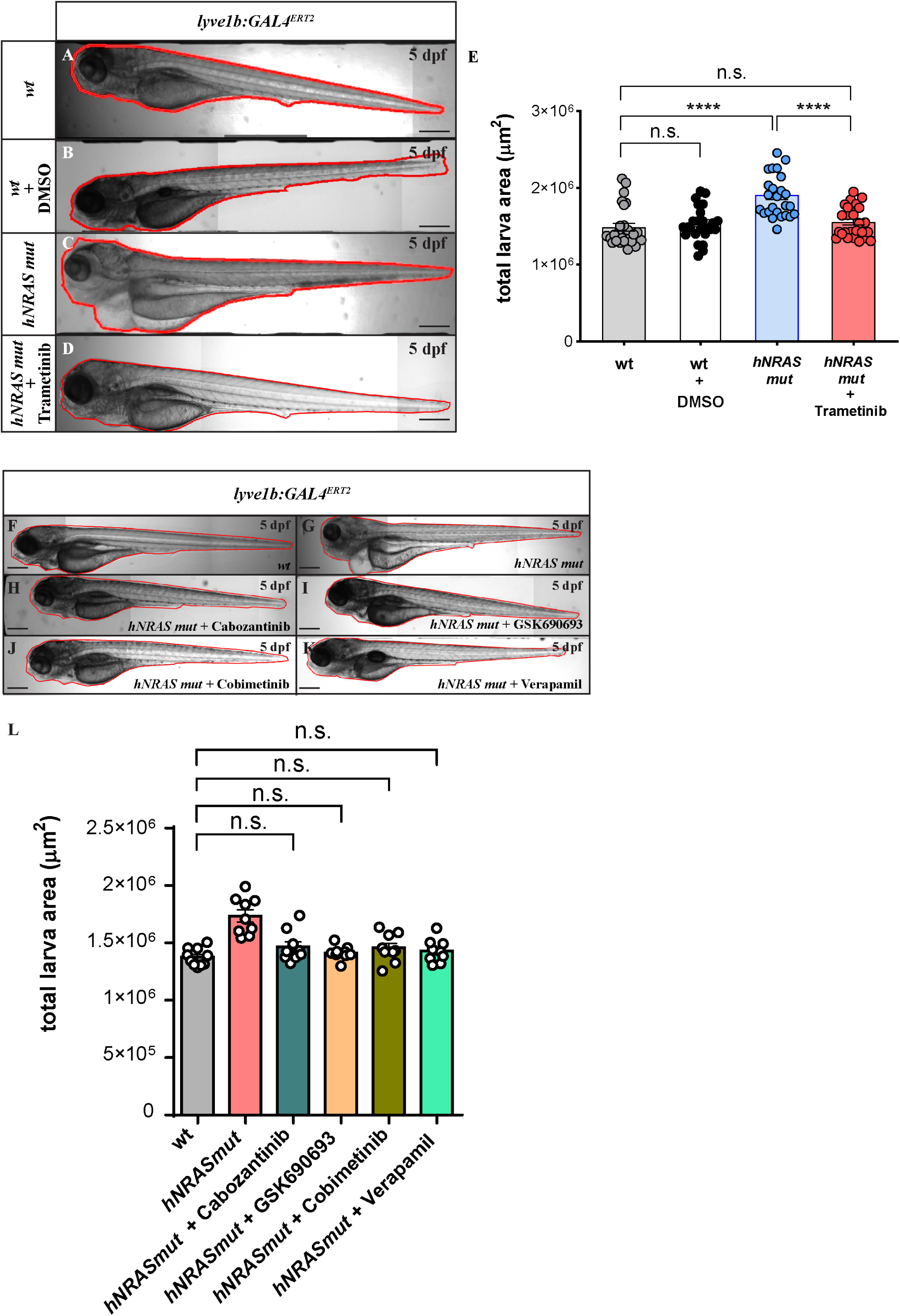
High throughput screen for small compounds reversing NRAS mutant phenotypes in zebrafish. **A-D.** Brightfield images of wt, wt + DMSO, *hNRASmut,* and *hNRASmut +* Trametinib 5 dpf *larvae*, obtained with WiScan® Hermes and processed by mask body segmentation with WiSoft® Athena Zebrafish Application. **E.** Quantification of total larva area in different samples (n=26; One-way ANOVA, multiple comparisons with Tukey post-hoc test). **F-L.** Brightfield images of 5 dpf wt (F), *hNRASmut* (G) and *hNRASmut* treated with Cabozantinib (H), GSK690693 (I), Cobimetinib (J) or Verapamil (K) showing complete rescue of total area in treated larvae (L) (n=10 One-way ANOVA, multiple comparisons with Tukey post-hoc test). Scale bars: A-D, F-J = 50 μm. Error bars are mean ± s.e.m.

For the first round of our screen, we compiled a library of 126 compounds, selected based on their association with known NRAS targets (Table 1, Fig. S2A), and added them at a concentration of 1uM. Out of this cohort, the algorithm automatically identified 35 compounds as significantly effective in reducing the total larva area to levels comparable to the wt group (Table 2). These hits were subjected to a second round of validation (Fig. S2B-E), by the end of which, we pinpointed 4 compounds that were most effective in reverting the “total embryo area” to average dimensions: GSK690693, Verapamil, Cabozantinib and Cobimetinib (Fig. 3F-L)

**Table 1.**
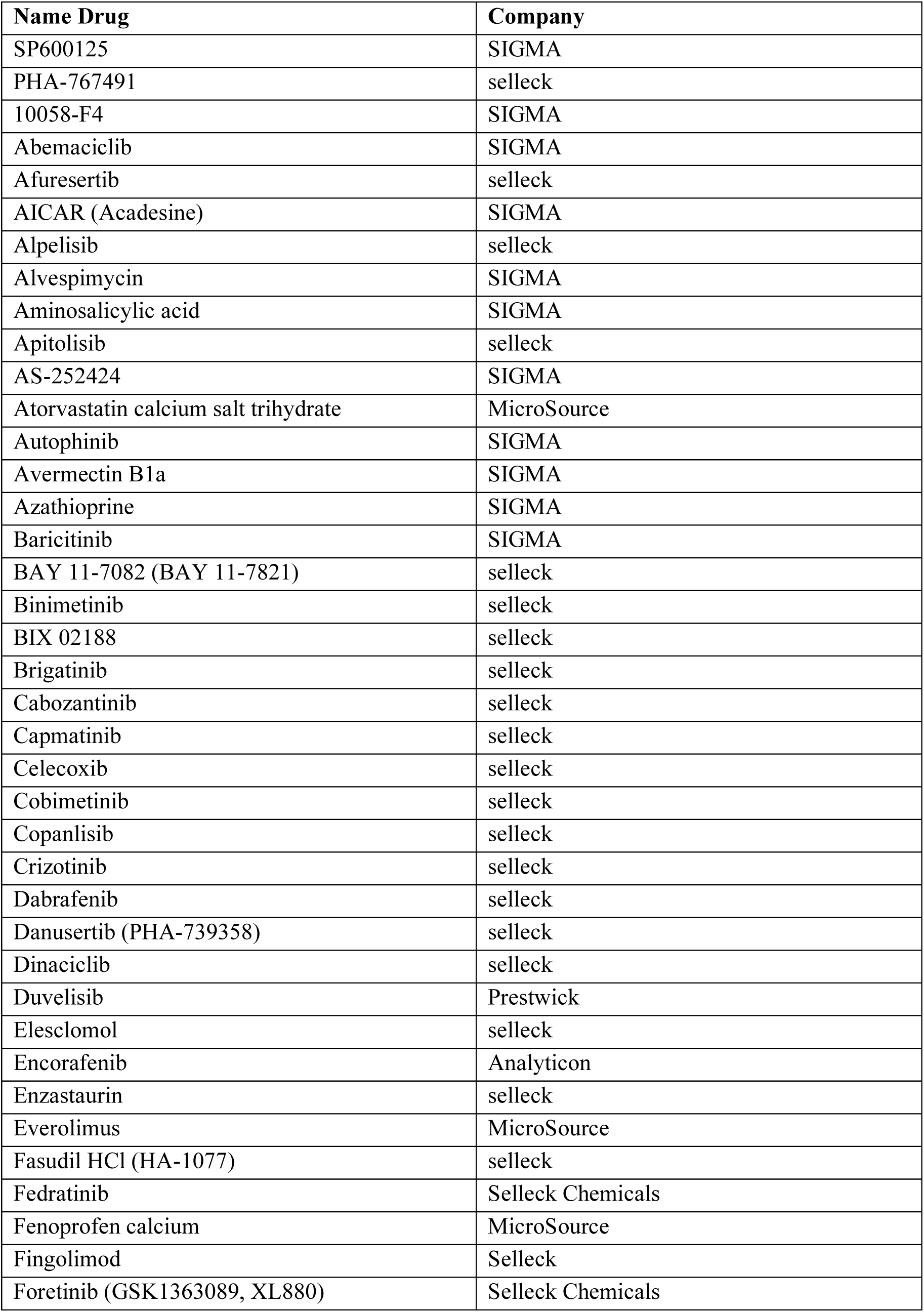
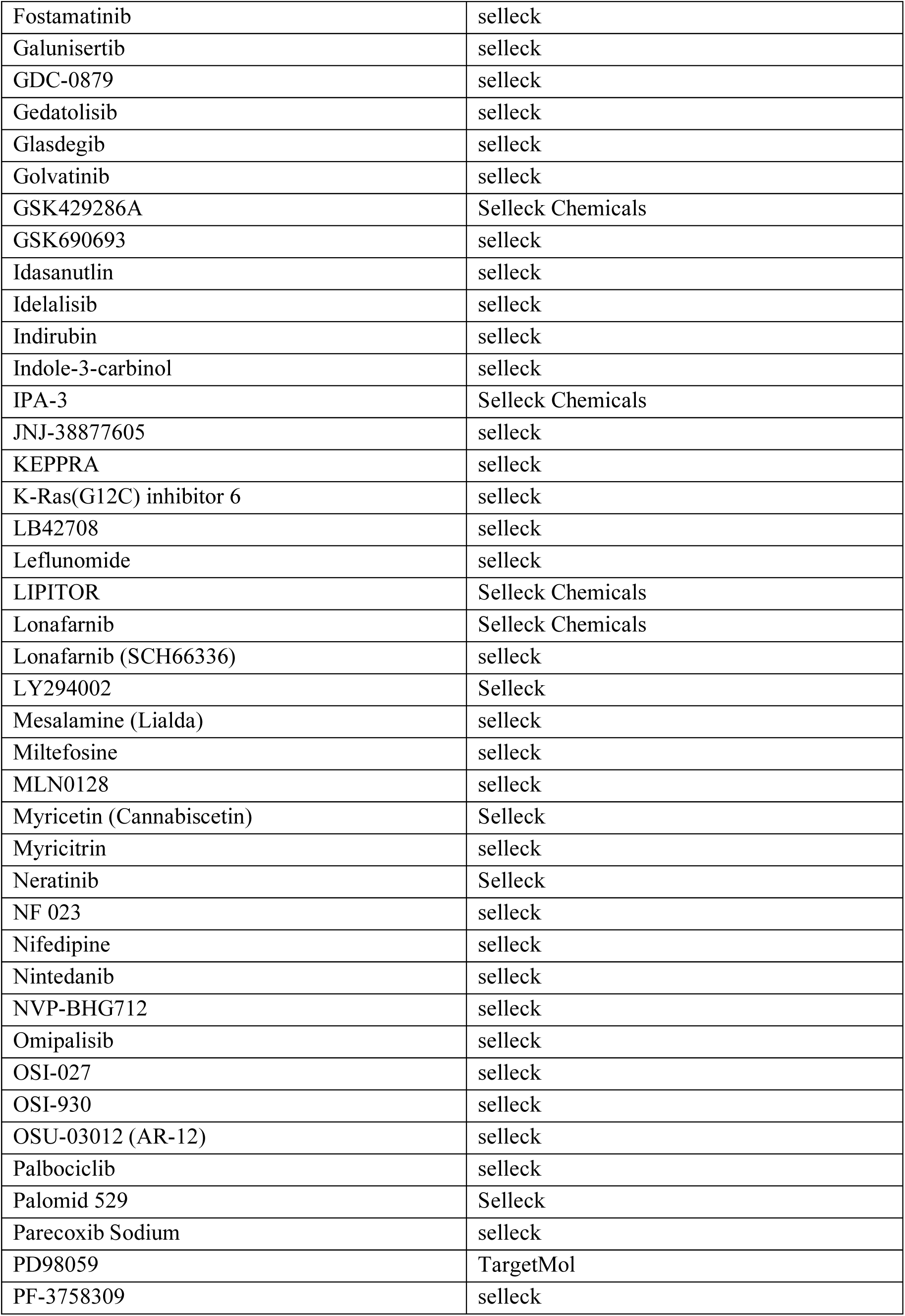
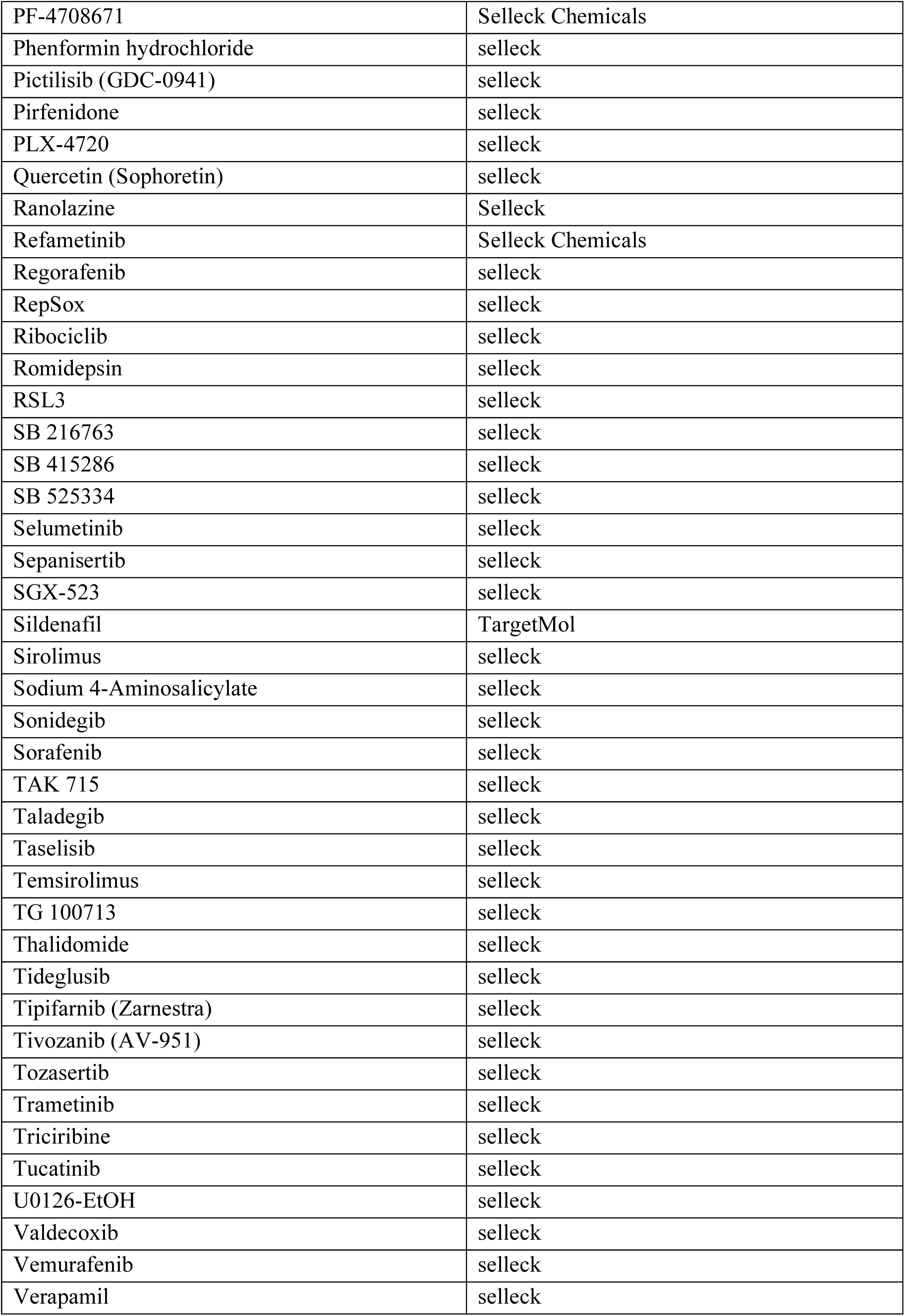
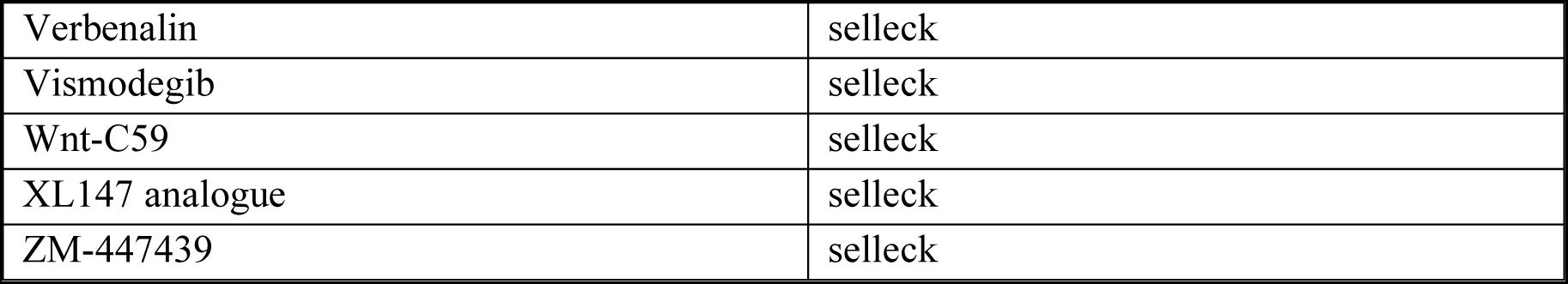
List of tested compounds.

**Table 2:**
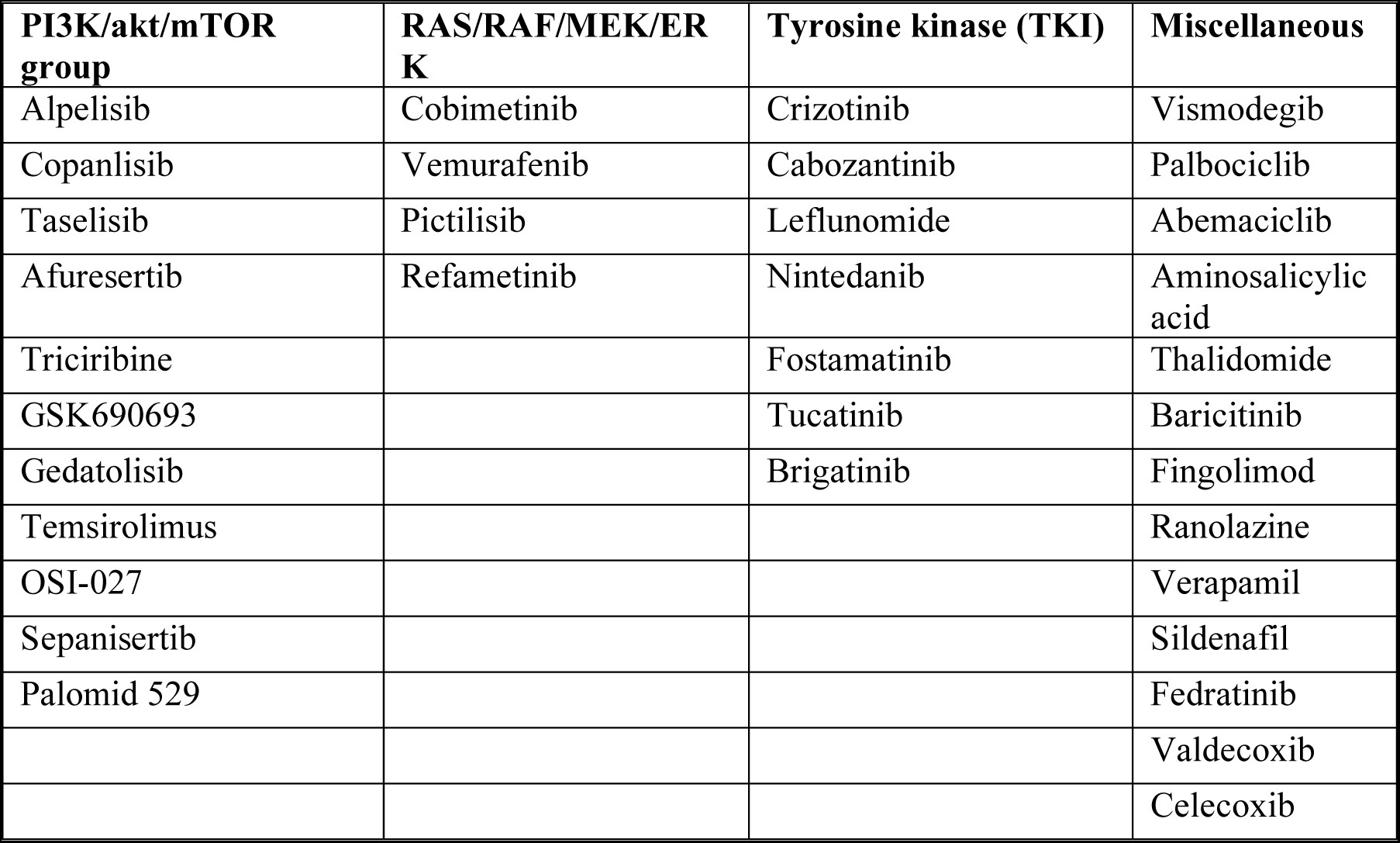
Selected compounds after high-throughput screening.

The next screening phase focused on determining the efficacy and specificity of the leading hits by analyzing TD formation after exposure to 3 different doses. Confocal images highlighted 3 compounds- GSK690693, Verapamil, and Cabozantinib- that were particularly effective in reverting the dilated TD phenotype of *hNRASmut* larvae, albeit showing slight differences in dose dependency (Fig. 4). GSK690693, a competitive pan-Akt inhibitor^32^ did not revert the TD dilation phenotype at concentrations of 0.5 and 1μM (Fig. 4C-E,G), as opposed to 2μM treatment that led to TD normalization (Fig.4F,G).

**Figure 4.**
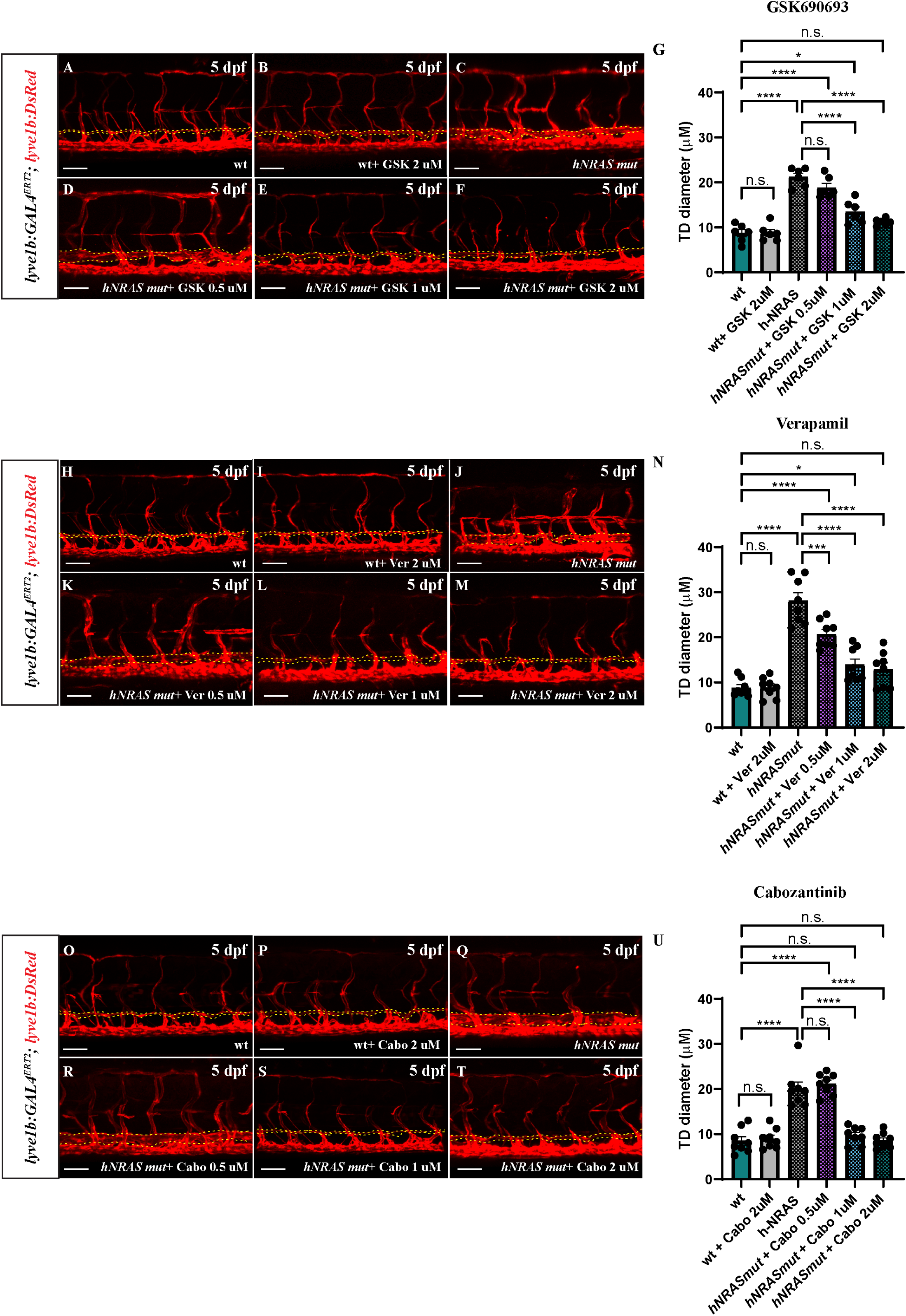
Selected candidates efficiently rescue the TD dilation phenotype in *hNRASmut larvae*. **A-G.** Confocal images of the trunk vasculature of 5 dpf *larvae* following treatment with Akt-Inhibitor GSK690693 (GSK) at 0.5 (D), 1 (E), and 2 (F) µM (quantified in G). **H-N.** Confocal images of the trunk vasculature of 5 dpf *larvae* following treatment with calcium channel-blocker Verapamil at 0.5 (K), 1 (L), and 2 (M) µM (quantified in N). **O-U.** Confocal images of the trunk vasculature of 5 dpf *larvae* following treatment with Cabozantinib at 0.5 (R), 1 (S), and 2 (T) µM (quantified in U). Yellow dashed lines in A-F, H-M, and O-T delineate the TD. Graphs G, N and U, n=6; One-way ANOVA, multiple comparisons with Tukey post-hoc test). Scale bars: A-F, H-M, O-T = 50 μm. Error bars are mean ± s.e.m.

Verapamil is a calcium-channel blocker used for the treatment of hypertension^33^. Embryos exposed to this compound showed a relatively mild effect at low dosages (Fig. 4K,N) and a complete rescue of TD caliber at the highest dose (Fig. 4L-N). We also assessed Cabozantinib, a receptor tyrosine kinase inhibitor with a broad range of targets, including MET, VEGFR2, FLT3, and c-KIT. Cabozantinib is currently FDA-approved for the treatment of medullary thyroid cancer, renal cell carcinoma, and hepatocellular carcinoma^34^. Like GSK690693, a 0.5µM dose was not effective, and embryos showed a TD diameter comparable with untreated *hNRASmut* samples (Fig. 4Q,R,U), while both 1μM and 2 μM resulted in a complete rescue of the phenotype (Fig. 4S-U). Finally, Cobimetinib failed to rescue the lymphatic phenotype based on confocal imaging analyses (Fig. S2F-L).

To verify the validity of the selected compounds as potential treatments for KLA, we tested their ability to inhibit LEC sprouting in NRAS p.Q61R mutant LECs isolated from the affected tissues of a KLA patient. We utilized a Spheroid-based sprouting assay, which recapitulates key features of sprouting lymphangiogenesis *in-vivo*^35^. Previous work demonstrated that the number of sprouts was increased in cultured KLA LECs^8^ as well as in Human Dermal LECs (HDLECs) transduced to express mutant NRAS p.Q61R^36^. As seen in Figure 5, we found that our screen hits Cabozantinib and GSK690693 were as effective as Trametinib in reducing sprout numbers (Fig. 5A-C,F). In addition, both elicited a significant decrease in p-S6 (Fig. 5G,H) without affecting pERK (Fig. 5I) levels. Interestingly, although Verapamil was effective in reversing the zebrafish phenotypes, it showed no impact on the sprouting of KLA cells (Fig. 5E) or p-S6 and p-ERK levels (Fig. 5G-I).

**Figure 5.**
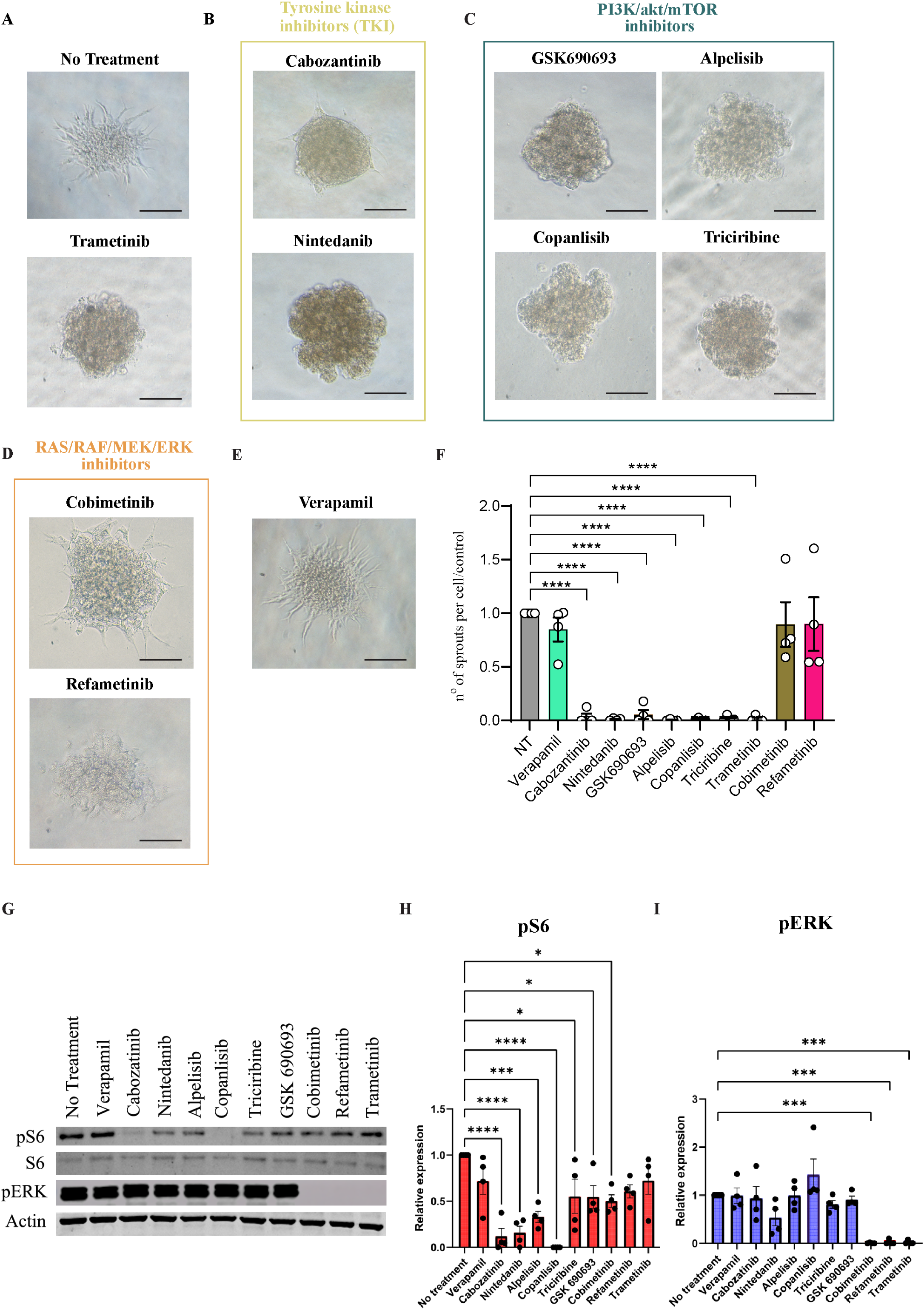
Validation in patient-derived cells. Representative images of sprout formation in KLA patient-derived LECs treated with the various drugs (2µM): **A.** no treatment (upper panel) Trametinib 0.5µM (bottom panel). **B.** Tyrosine Kinase inhibitors. **C.** P13K/Akt/mTOR inhibitor. **D.** RAS/RAF/MEK/ERK inhibitors. **E.** Verapamil. **F.** Quantification of numbers of sprouts calculated from four independent experiments (5 wells, ∼200 spheroids per condition). Plotted are means ± s.e.m., * indicates p < 0.05. **G-I**. WB analysis of the effect of drugs (at 2µM) on phosphorylated ERK (p-ERK) and phosphorylated S6 (p-S6). Plotted are means ± s.e.m., * indicates p < 0.05. n=4; One-way ANOVA. Scale bar: A-E= 100μm

We next asked whether the positive effects of GSK690693 and Cabozantinib were specific to these compounds or more broadly common to other drugs belonging to the same family and inhibiting the same pathways. Indeed, Nintedanib, a TKI targeting vascular endothelial growth factor receptor (VEGFR) 1, 2, and 3 among other RTKs^37^ exerted a significant inhibitory effect on LEC sprouting (Fig. 5B) and p-S6 levels (Fig. 5G,H, similar to Cabozantinib). To validate and further expand the positive results exerted by GSK690693 on PI3K-AKT-mTOR, we tested Alpelisib (PI3K inhibitor), Copanlisib (PI3K inhibitor), and Triciribine (AKT inhibitor), all of which inhibited LEC sprouting and p-S6 at comparable levels. Interestingly, 2 MEK inhibitors (Cobimetinib and Refametinib) did not inhibit sprouting, albeit suppressing ERK phosphorylation.

Finally, we stained the KLA patient’s tissues with pAKT and pERK antibodies to get insight into the activation of these pathways *in situ*. Appendix served as a positive control for Podoplanin, and colon carcinoma as a positive control for pAKT^38^ and pERK^39^ staining. As seen in Fig S3A-B, both pAKT and pERK are detected in KLA abnormal lymphatic vessels. However, pAKT levels are more specific and restricted to the Podoplanin+ lymphatics.

Taken together, our results demonstrate the suitability of our novel zebrafish model for studying the pathophysiology of KLA and identifying novel avenues of treatment. Moreover, the combination of the zebrafish and human assays highlights tyrosine kinase-, as well as PI3K-AKT inhibition, as potential pharmaceutical avenues for treating KLA.

## Discussion

To model KLA in zebrafish we established two transgenic lines-*prox1a:KALTA4;hNRASmut/wt* constitutively expressing the mutant or wt forms of human NRAS under the control of the *prox1a* promoter, and a line based on the GAL4-UAS system, enabling spatiotemporal control of *hNRASmut/wt* expression under the *lyve1b* promoter and controlled by the estrogen receptor ERT2. We found that both lines recapitulated clinical features of KLA, including pericardial edema and a dilated TD, albeit expression driven by the *prox1a* promoter resulted in general toxicity and non-specific effects primarily due to the early and broad expression of the *prox1a* gene. We further found that Trametinib, a MEK inhibitor currently used in the clinic for KLA treatment, was highly efficient in reversing the mutant phenotypes. Finally, our AI-based screening of 126 compounds identified GSK690693, a pan-AKT inhibitor, and Cabozantinib, a tyrosine kinase inhibitor, as novel drug candidates for treating KLA based on their ability to revert the zebrafish lymphatic phenotype, normalize patient’s LEC function, and block phosphorylation of NRAS downstream targets.

Interestingly, Verapamil ameliorated the zebrafish phenotype in a dose-dependent manner but did not hinder the patient’s abnormal LEC sprouting or downstream signaling through pAKT, pERK, and pS6. Calcium signaling can regulate Ras activity (NRAS, HRAS, and KRAS) by numerous mechanisms^40^. Moreover, RAS oncogenic mutations were shown to influence cytosolic Ca2+ signals^41,42^. However, these reciprocal effects are cell and context-dependent^40^. Accordingly, Verapamil’s effect could be limited to a subpopulation of lymphatic progenitors existing in the zebrafish mutant, which is not present in the human-cell assay. Alternatively, Calcium modulation in NRAS-mutated LECs might couple to different Ras effectors, other than the ERK/MAPK cascade, such as Ral/GDS and PLCε^40^, and affect functions that are not recapitulated by the sprouting assay. Future clinical studies are needed to verify whether Verapamil can ameliorate KLA phenotype as a monotherapy or in combination with other drugs such as Trametinib.

GSK690693 is an ATP-competitive pan-Akt kinase inhibitor. To date, several allosteric and ATP-competitive AKT inhibitors have been synthesized and tested in clinical trials for various cancers^43,44^. Unlike tumors that often exhibit numerous somatic mutations, vascular anomalies, including KLA, have been demonstrated to be somatic monogenic disorders. Consequently, they are more likely to rely on the presence of a driver oncoprotein for their development and progression. Indeed, AKT inhibition with Miransertib was shown to improve the phenotype of Proteus syndrome^45^, an overgrowth disorder caused by postzygotic activating variants in AKT1 and two children with PIK3CA-related overgrowth spectrum (PROS), with a favorable safety profile^46^.

The efficiency of GSK690693 in reverting the KLA phenotype reaffirms the critical role of the PI3K/AKT pathway as RAS effector during lymphatic development and is in line with previous findings on the role of RAS/PI3K interactions in a mouse model field^47^. By generating a mouse in which the binding of RAS to the p110α subunit of PI3K was prevented by the introduction of two point mutations in the RBD domain of the *Pik3ca* gene, it was shown that homozygous mutant mice died shortly after birth due to chylous ascites resulting from deficient lymphatic development.

Cabozantinib is a tyrosine kinase inhibitor, FDA-approved for treating several malignancies, including in pediatric patients^34,48^. Its targets include VEGF receptors 1, 2, and 3^49^, further supporting its effect on lymphangiogenesis. Although adverse effects are common, they are dose-related, leading to treatment discontinuation in only 9–20% of patients.

Interestingly, mTOR inhibitors had only a partial effect on the abnormal lymphatic phenotype in our zebrafish model. These findings are in accordance with clinical data, as no complete response was obtained in patients, and only half of KLA patients achieved a partial response^36^.

Our findings reaffirm the previously identified role of the Q61R NRAS mutation in the lymphatic endothelium in the pathogenesis of KLA. Indeed, this mutation was identified in the involved tissue and body fluids of most KLA patients^50^. However, the timing of appearance of the mutation remains largely unknown. The young presentation age of most patients and the multifocal nature of the disease may suggest that the mutation occurs early, during embryonic development. Our findings of a specific "time window" required for the mutation to exert a phenotype might support this hypothesis.

There are some limitations to our use of the zebrafish model. First, as microscopic imaging of adult zebrafish remains highly challenging, we focus on early embryo/larval stages. Thus, it remains to be tested in other animal models to determine whether similar efficacy of the drugs is observed in adult animals. Moreover, since drug administration in zebrafish is done preferentially through addition to the embryo water, drug screening for insoluble drugs remains challenging in this model. Finally, the physiology of zebrafish presents evolutionary divergences compared to mammals, such as optimum temperature and breath system that can affect drug absorption and distribution, enhancing the importance of further characterizing the pharmacokinetics in mammalian systems. Nevertheless, our results demonstrating that 2 of the lead candidates identified in the zebrafish-based screen successfully reversed the sprouting phenotype of KLA cells strongly emphasize the relevance of this model for preclinical drug assessment of novel therapeutic avenues for KLA treatment.

## Acknowledgments

The authors thank H. Hasid and Y. Yogev (Weizmann Institute, Israel) for technical assistance; G. Almog, R. Hofi, A. Glozman, and R. Brihon (Weizmann Institute, Israel) for superb fish care. A. Pavlovsky (Inst. of Pathology, Sheba Medical Center) for assistance with immunostaining. G. Cohen (Wohl Institute for Drug Discovery) for compound plate preparation; E. Ben Zeev (Medicinal Chemistry, G-INCPM) for chemical database searches. This work was supported in part by ERC CoG (LymphMap 818858) to KY, Million Dollar Bike Ride Grant (MDBR-23-021-CLA) from the Orphan Disease Center, University of Pennsylvania to KY, the Weizmann SABRA – Yeda-Sela – WRC Program to KY, the Estate of Emile Mimran, and The Maurice and Vivienne Wohl Biology Endowment and the Brenden-Mann Women’s Innovation Impact Fund to KY, Lymphatic Malformation Institute grant to SG, Orphan Disease Center 2018 Million Dollar Bike Ride to SG and Talpiot Medical Leadership grant, Sheba Medical Center, to SG. K.Y. is the incumbent of the Enid Barden and Aaron J. Jade Professorial Chair in Memory of Cantor John Y. Jade. I.B. was supported by a postdoctoral fellowship by the Sergio Lombroso Program and a Senior Postdoc Fellowship by the Weizmann Institute. N.M. is supported by research grants from the Estate of Olga Klein Astrachan and the Estate of Mady Dukler.

## Author contributions

I.B. designed and conducted experiments, analyzed data, and co-wrote the manuscript; A.J., S.E., N.F., J.L. conducted experiments and data analyses; G.L designed and conducted experiments, and analyzed data; N.F, P.G, L.L, A.B, and K.A conducted experiments and data analyses; J.J.O. and Y.P. developed and adapted the WiScan® Hermes automated microscope and the WiSoft® Athena software and assisted with drug screen; N.M. assisted with fish experiments; H.B. devised and assisted with drug screen; K.Y. and S.G. directed the study, secured funding, designed experiments, analyzed data and co-wrote the paper with inputs from all authors.

Y.P and J.J.O. are employed by IDEA Bio-Medical. Other authors declare no competing interests.

## Materials and Methods

### Zebrafish husbandry, transgenic and mutant lines

Zebrafish were raised by standard methods and handled according to the guidelines of the Weizmann Institute Animal Care and Use Committee^23^. For all imaging, embryos were treated with 0.003% phenylthiourea (PTU, Sigma-Aldrich) from 8 hpf to inhibit pigment formation. Zebrafish lines used in this study were: *Tg(fli1:EGFP)^yl^* ^51^*; Tg(kdrl:TagBFP)^mu2^*^93^ ^25^; *TgBAC(prox1a:KALTA4,4xUAS-E1B:TagRFP) ^nim5^* ^24^.

The *Tg(lyve1b:Gal4^ERT2^)* construct was generated by combining the *p5E-lyve1b* promoter^52^ with the ERT2 sequence and then with *pME-Gal4VP16*^53^ using the Gateway System.

The *Tg(UAS:hNRASwt)* and *Tg(UAS:hNRASmut)* were generated by cloning the human wt and p.Q61R NRAS variants, respectively, into Tol2-compatible vectors. The resulting constructs were injected at the 1-cell stage into AB zebrafish to generate stable lines ^54^.

### Imaging and image processing

Confocal imaging was performed using the Zeiss LSM780 or LSM880 upright confocal microscopes (Carl Zeiss) equipped with a water-immersed ×20/1.0 NA or ×10/0.5 NA objective lens. Embryos were mounted using 1.5% (w/v) low-melting agarose. Confocal images were processed offline using the Fiji^55^ version of ImageJ (NIH) or Imaris v.9.3 (Bitplane). The fluorescent images shown in this study are single-views, 2D-reconstructions, of collected z-series stacks.

Phase Contrast images of zebrafish embryos were obtained using a stereomicroscope LEICA M165 FC equipped with a Nikon DS-Fi3.

For drug screening, embryos were incubated in transparent glass bottom 96-well plates (ZF plates, Hashimoto, Mie, Japan). Transmitted light images were obtained using WiScan® Hermes (IDEA Bio-Medical, Rehovot, Israel), a high-content inverted scanning system equipped with 4X air objectives with high numerical aperture (NA)^31^. Images of each well were stitched to form a single image of the full larva body.

Drug screening image analysis was performed using the WiSoft® Athena Software Zebrafish Application (IDEA Bio-Medical, Rehovot, Israel). Athena utilizes a novel deep learning-based AI algorithm to analyze brightfield images of the whole zebrafish larvae to identify the fish outline and certain anatomical structures, such as heart, eye, tail, and three body compartments (head, trunk, and tail) as previously described^31^. Eye and tail anatomical structures were utilized to select only properly oriented fish to permit total larva area comparisons. In the assay development process, a manual drawing tool embedded in the WiSoft® Athena software was used to adjust mask definitions.

### Tamoxifen induction and drug screen

*Tg(lyve1b:Gal4^ERT2^;UAS:hNRASwt)* and *Tg(lyve1b:Gal4^ERT2^; UAS:hNRASmut) larvae* were raised until 24 hpf when 5 µM of Tamoxifen was added. At 48 hpf, Tamoxifen was washed, and inhibitors were added. To perform a focused, clinically relevant screen, we first defined 126 canonical and non-canonical NRAS targets based on current literature (Fig. S3A,B). The Drug Bank database was used for mining compounds with NRAS targets as a list of keywords. We then prioritized the compounds based on their approval state (clinically approved compounds were given priority) and level of association with NRAS targets based on target annotations from the supplier. Known cytotoxic drugs CPM were excluded. High-throughput drug screening was conducted in 96-well plates with a well-slit design that allows efficient alignment of zebrafish and optical clear bottom for imaging (ZF plates, Hashimoto, Mie, Japan). Compounds for screening were picked from source plates and transferred to assay plates using an Echo 555 acoustic transfer liquid handler (Labcyte/Beckman). Each drug was tested at a concentration of 1 µM directly added in the Hashimoto plate, and then fish were kept in dark conditions in a 28 ⁰C incubator until day 5 when they were imaged using the WiScan® Hermes (IDEA Bio-Medical, Rehovot, Israel).

### Western blotting

Isolation and characterization of KLA patient Lymphatic endothelial cells were previously derscribed^8^. The study was approved by the institutional research committee at Sheba Medical Center (#8333-10-SMC). Cells were seeded at 60% confluence in 60X15 mm dishes. 24 hours later, the cells were either not treated or treated with 1μM of one of the following drugs: Verapamil, Cabozantinib, Nintedanib, Alpelisib, Copanlisib, Triciribine, GSK-690693, Cobimetinib, Refametinib, Trametinib. Following a 24-hour incubation, whole cell lysates were generated. The cells were washed twice with PBS and scraped with a cell scraper in RIPA lysis buffer system (Cat. No. sc-24948, Santa Crus Biotechnology, CA, USA). The cells were incubated with the buffer on ice for 30 minutes, and then cellular debris was removed by centrifugation at 14,000x RPM for 30 minutes at 4°C. Protein concentration was determined by Micro BCA^TM^ Protein Assay Kit (ThermoFisherscientific, MA, USA) according to the manufacturer’s instructions. 30 μg protein of each cell lysate were separated by 4-20% gradient Bis-Tris- PAGE (SurePAGE^TM^, Bis-Tris, Cat. No: M00657, A2S technologies, Israel) in Tris-MOPS-SDS running buffer (Cat. No: M00138, A2S technologies, Israel) followed by transfer to nitrocellulose sheets. Then, the membranes were blocked for 1 hour in Intercept Blocking Buffer (Cat. No: 927-60001, LI-COR, NE, USA). Next, the membranes were incubated with primary antibodies overnight at 4°C. After washing three times for 10 minutes with TBST (Cat. No 002089232300, Bio-Lab, Israel) diluted 1 to 10 in distilled water, the membranes were incubated with the appropriate goat anti-mouse or goat anti-rabbit IgG peroxidase conjugate for 1 hour at room temperature. Next, the membranes were rewashed with TBST as described, subjected to ECL substrate (WESTAR ANTARES, Cat. No. XLS142,0250, Cyanagen, Italy), and the signal was detected using the ChemiDOc^TM^ MP Imaging System (Bio-Rad, CA, USA). Finally, densitometric analysis of the detected bands was performed using Image Lab software (Bio-Rad, CA, USA). Primary antibodies used: Monoclonal rabbit anti-Phospho- p44/42 MAPK (Erk1/2) (Thr202/Tyr204) (20G11) (Cat. No. 4376S; Monoclonal rabbit anti-S6 Ribosomal Protein (5G10) (Cat. No. 2217); Monoclonal rabbit anti-Phospho-S6 Ribosomal Protein (Ser235/236) (2F9) (Cat. No. 4856); (All were purchased from Cell Signaling Technology Inc, MA, USA) {dilution 1:1000}. Monoclonal mouse anti-β-Actin (AC-15) (Cat. No. sc-69879); (All were purchased from Santa Crus Biotechnology, CA, USA) {dilution 1:1000}

### LEC sprout assay

The isolation and characterization of KLA Patient-derived LEC were previously described^8^. 6X10^5^ cells were plated in 24-well dishes AggreWell400 pretreated with AggreWell Rising solution (STEMCELL Technologies) in EBM2 medium for 16 h to allow the formation of spheroids containing ∼ 500 cells. On day 2, the spheroids were collected and filtered through a 40-µm pluriStrainer (pluriSelect) to ensure uniform spheroid size. The spheroid yield was quantified by microscopy, and 8-10 spheroids were mixed with Cultrex Basement Membrane Extract (Trevigen) to a final protein concertation of 8 mg\ml with the addition of the various treatments and embedded within a single well of a 96-well plate. The next day, for each condition, five wells corresponding to ∼50 spheroids in total, were manually photographed using a Nikon Eclipse TS-100 microscope (10x objective) and a Nikon DS-Fi1 camera, and the numbers of sprouts were manually counted.

### Immunostaining

FFPE blocks from KLA-infiltrated tissues^8^ were sectioned at 4 μm and a positive control was added on the right edge of the slides. Phospho-AKT (#9271, Cell Signaling, USA), phosphor-ERK1/2 (#4376, Cell Signaling, USA), and Podoplanin (M3619, Dako, USA) immunostaining protocols were calibrated on a Benchmark Ultra staining module (Ventana Medical Systems Inc., USA). Slides were warmed to 60°C for 1 hour and then processed by a fully automated protocol. Briefly, after dewaxing and rehydration, sections were exposed to 64-minute CC1 HIER pretreatment (Ventana Medical Systems Inc., USA) for pAKT and pERK1/2. Sections were pretreated for 36 minutes for Podoplanin. pAKT antibody (1:25), pERK1/2 antibody (1:50), and Podoplanin (1:80) were all incubated at 37°C for 44 minutes. PAKT and pERK1/2 were detected with an OptiView detection kit (Ventana Medical Systems). pAKT staining was enhanced with the OptiView Amplification Kit (Ventana Medical Systems Inc., USA). Podoplanin was detected with an UltraView detection kit (Ventana Medical Systems). Afterward, all sections were counterstained with hematoxylin (Ventana Medical Systems). At the end of the automated run, slides were dehydrated in graded ethanol (70%, 96%, and 100%). Before cover-slipping, sections were cleared in Xylene and mounted with Entellan.

## Statistical analyses

Comparison of two samples was done using unpaired two-tailed Student’s t-test assuming equal variances from at least three independent experiments unless stated otherwise. Statistical significance for three or more samples was calculated via one-way ANOVA followed by Tukey’s or Dunnett’s multiple comparisons test unless otherwise indicated. All data are reported as mean values ± SEM and were analyzed using Prism 6 software (GraphPad Software, Incorporated, La Jolla, CA, USA).

---

**Figure S1.**
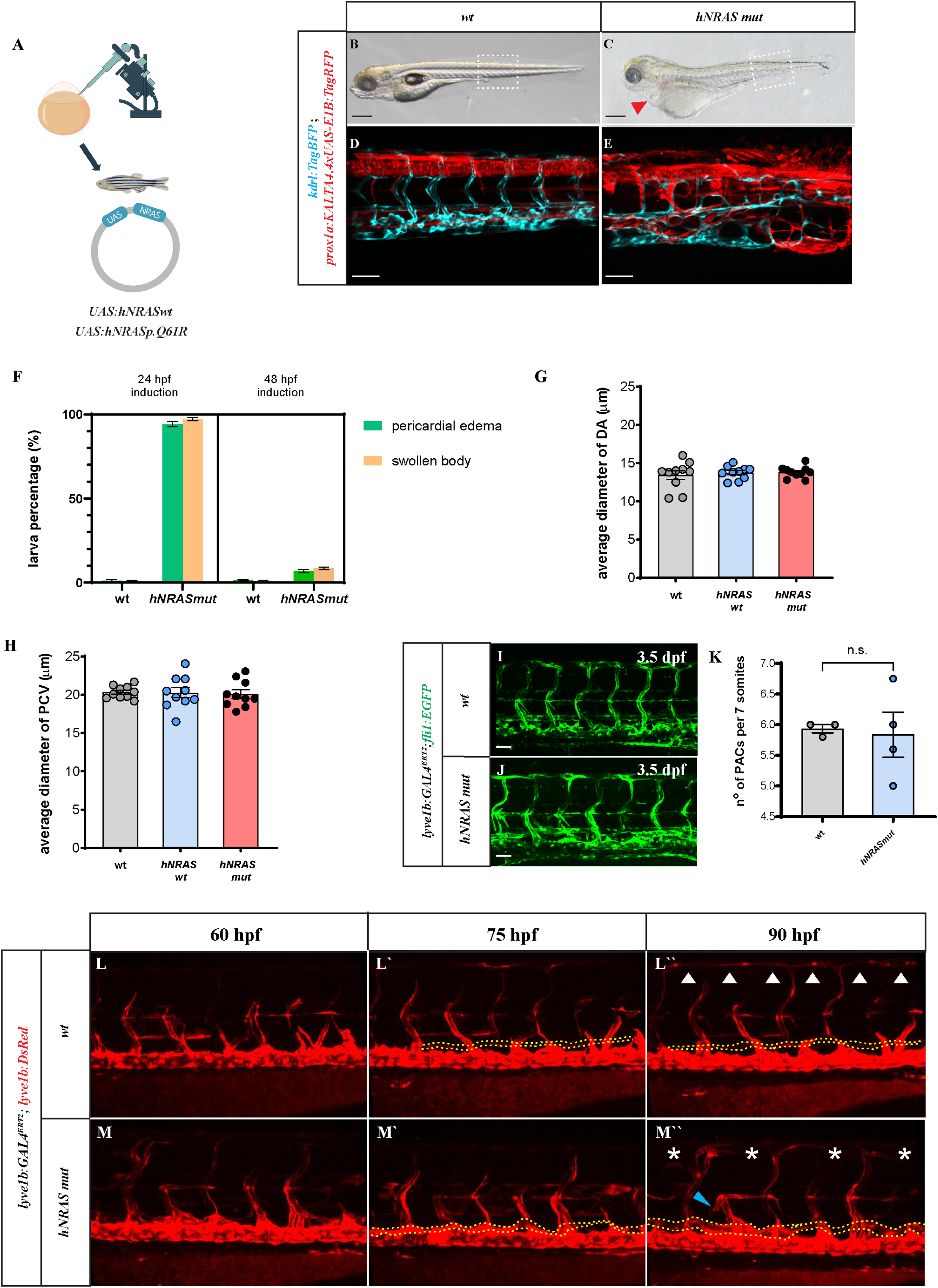
Generation of Tg*(UAS:hNRAS)* zebrafish transgenic lines. **A.** Diagram depicting the strategy used to establish Tg*(UAS:hNRASwt)* and Tg*(UAS:hNRASmut)* transgenic lines. **B-E.** Constitutive expression of *hNRASmut* under *prox1a* promoter results in pericardial edema, swollen body (C), severely malformed trunk vasculature, and general toxicity (E). Dashed squares in B-C mark the area enlarged in D,E). **F.** Quantification of morphological defects following induction of *hNRASmut* expression in *lyve1b:GAL4^ERT2^*embryos, at 24 and 48 hpf (N=3, n=20). **G,H.** Quantification of DA and the PCV caliber after induction of *hNRASwt* and *hNRASmut* in *lyve1b:GAL4^ERT2^*embryos (n=10, One-way ANOVA, multiple comparisons with Tukey post-hoc test). **I-K.** Confocal images of the trunk of wt (I) and *hNRASmut* (J) embryos at 3.5 dpf showing no significant differences in the number of PACs, quantified in (K, n=4, two-tailed Student’s t-test, *P*=0.83). **L-M’’.** Selected images from confocal time-lapse series of *Tg(fli1:EGFP; lyve1b:DsRed; lyve1b:GAL4^ERT2^)* wt (L-L’’) or following induction of *hNRASmut* expression (M-M’’) showing dilated PAC sprouts (M’’, blue arrowheads), bloated TD (M’’, yellow dashed line) and missing DLLV (M’’, asteriks); white arrowheads in L’’ point to normal DLLV. Scale bars: B,C = 100 μm; D,E = 100 μm; J,K= 50 μm; L-M’’ = 100 μm. Error bars are mean ± s.e.m.

**Figure S2.**
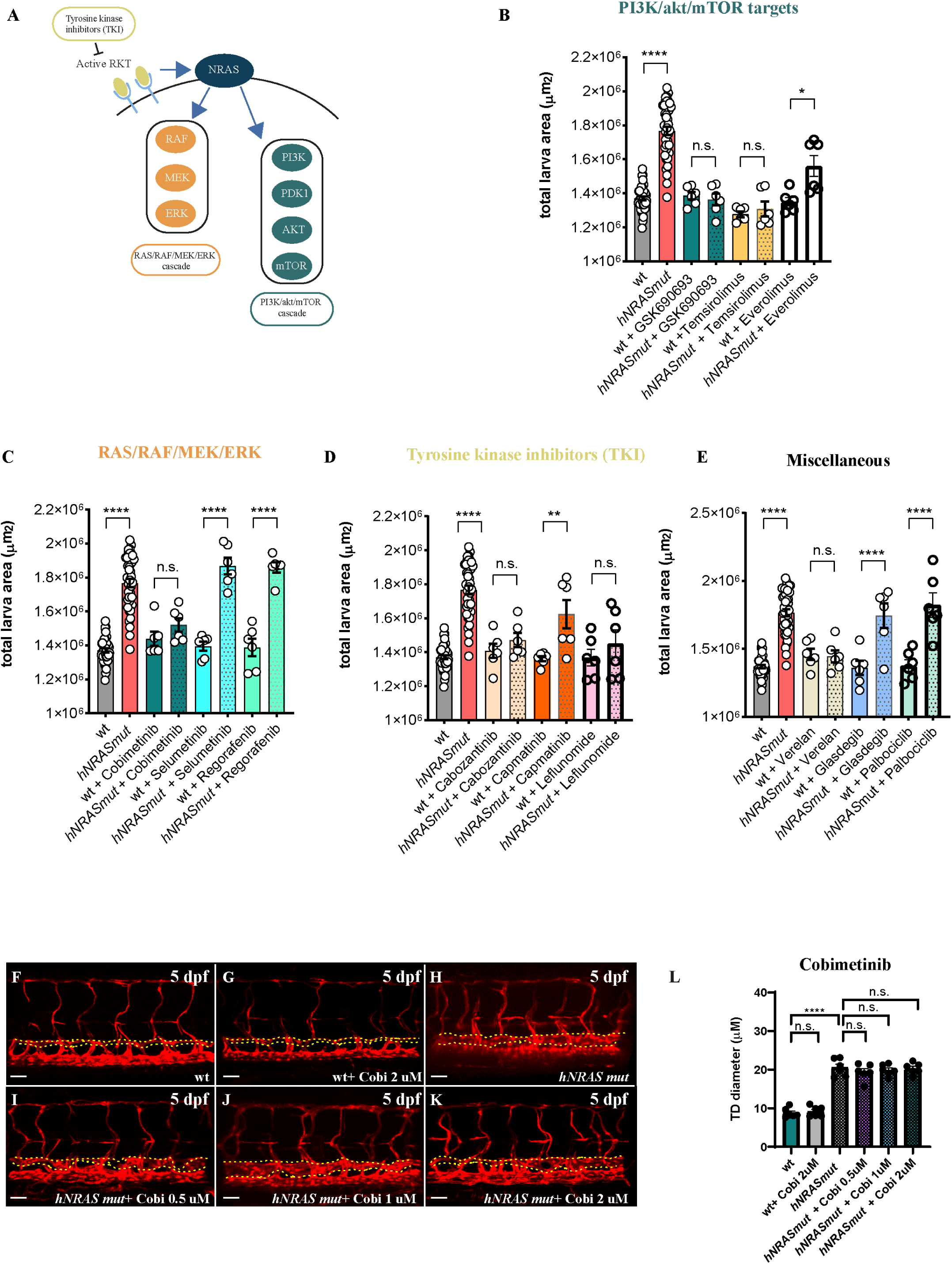
High-throughput drug screening in zebrafish larvae. **A.** Diagram indicating NRAS downstream targets selected for screening. **B-E**.Quantification of the total fish area following segmentation and analysis with WiSoft® Athena Zebrafish Application. Representative compounds from each target group are shown (n_wt_=40, n*_hNRASmut_*=40, n_wt+single compound_=6 and n*_hNRASmut_*_+single compound_=6; One-way ANOVA, multiple comparisons with Tukey post-hoc test). **F-L.** Confocal images of trunk vasculature following Cobimetinib administration show no rescue of TD dilation phenotype (TD highlighted in yellow (n=6; One-way ANOVA, multiple comparisons with Tukey post-hoc test). Error bars are mean ± s.e.m. Scale bars: F-L = 50 μm

**Figure S3.**
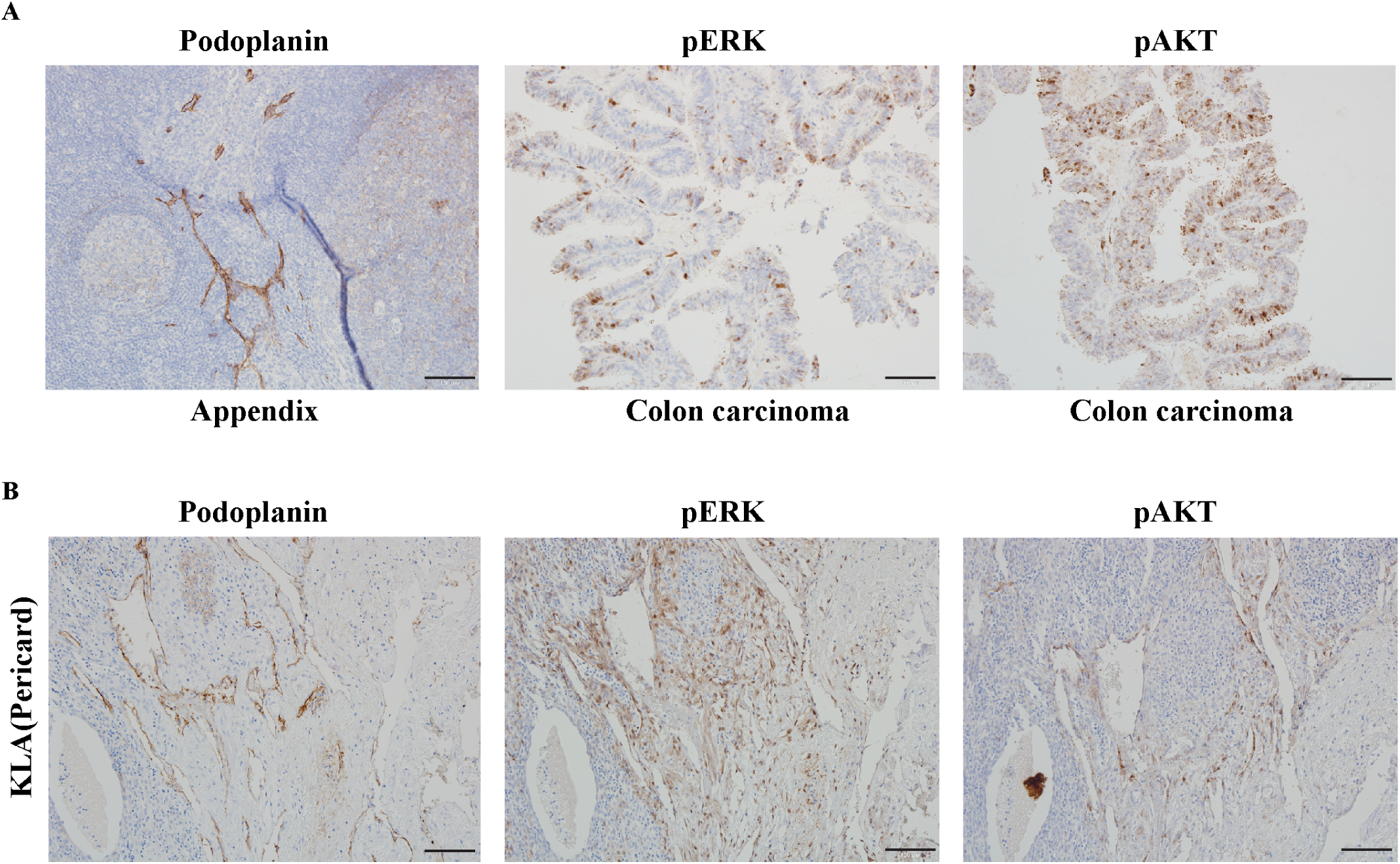
pAKT and pERK levels are increased in KLA lymphatic vessels. **A**. Positive controls for Podoplanin, pERK and pAKT staining. **B**. Staining of patient’s KLA involved tissue for Podoplanin, pERK and pAKT staining; H = 100μm

**Video S1**: Time-lapse series of *Tg(lyve1b:GAL4^ERT2^; lyve1b:DsRed)* wt (left) and *hNRASmut* (right) embryos showing trunk lymphatic development. White arrowhead points to defective PAC.

